# Gα/GSA-1 works upstream of PKA/KIN-1 to regulate calcium signaling and contractility in the *Caenorhabditis elegans* spermatheca

**DOI:** 10.1101/2020.02.03.932871

**Authors:** Perla G. Castaneda, Alyssa D. Cecchetelli, Hannah N. Pettit, Erin J. Cram

**Affiliations:** Department of Biology, Northeastern University, Boston, MA 02115, United States

## Abstract

Correct regulation of cell contractility is critical for the function of many biological systems. The reproductive system of the hermaphroditic nematode *C. elegans* contains a contractile tube of myoepithelial cells known as the spermatheca, which stores sperm and is the site of oocyte fertilization. Regulated contraction of the spermatheca pushes the embryo into the uterus. Cell contractility in the spermatheca is dependent on actin and myosin and is regulated, in part, by Ca^2+^ signaling through the phospholipase PLC-1, which mediates Ca^2+^ release from the endoplasmic reticulum. Here, we describe a novel role for GSA-1/Gα_s_, and protein kinase A, composed of the catalytic subunit KIN-1/PKA-C and the regulatory subunit KIN-2/PKA-R, in the regulation of Ca^2+^ release and contractility in the *C. elegans* spermatheca. Without GSA-1/Gα_s_ or KIN-1/PKA-C, Ca^2+^ is not released, and oocytes become trapped in the spermatheca. Conversely, when PKA is activated through either a gain of function allele in GSA-1 (GSA-1(GF)) or by depletion of KIN-2/PKA-R, Ca^2+^ is increased, and waves of Ca^2+^ travel across the spermatheca even in the absence of oocyte entry. In the spermathecal-uterine valve, loss of GSA-1/Gα_s_ or KIN-1/PKA-C results in sustained, high levels of Ca^2+^ and a loss of coordination between the spermathecal bag and sp-ut valve. Additionally, we show that depleting phosphodiesterase PDE-6 levels alters contractility and Ca^2+^ dynamics in the spermatheca, and that the GPB-1 and GPB-2 G_β_ subunits play a central role in regulating spermathecal contractility and Ca^2+^ signaling. This work identifies a signaling network in which Ca^2+^ and cAMP pathways work together to coordinate spermathecal contractility.

**Author Summary:** Organisms are full of biological tubes that transport substances such as food, liquids, and air through the body. Moving these substances in a coordinated manner, with the correct directionality, timing, and rate is critical for organism health. In this study we used *Caenorhabditis elegans*, a small transparent worm, to study how cells in biological tubes coordinate how and when they squeeze and relax. The *C. elegans* spermatheca is part of the reproductive system, which uses calcium signaling to drive the coordinated contractions that push fertilized eggs out into the uterus. Using genetic analysis and a calcium-sensitive fluorescent protein, we show that the G-protein GSA-1 functions with protein kinase A to regulate calcium release, and contraction of the spermatheca. These findings establish a link between G-protein and cAMP signaling that may apply to similar signaling pathways in other systems.

## Introduction

Regulation of cellular contractility and relaxation is essential for the function of epithelial and endothelial tubes, which are subjected to changing flow, strain, and pressure as they transport liquid, gases, and other cells throughout the body [1,2]. The *Caenorhabditis elegans* spermatheca, part of the hermaphroditic reproductive system, is an excellent model for the study of coordinated cell contractility [3–8]. The *C. elegans* reproductive system is composed of two symmetrical gonad arms surrounded by smooth muscle-like sheath cells, the spermathecae, and a common uterus [9,10]. The spermatheca, the site of fertilization, consists of an 8-cell distal neck, a 16-cell central bag, and a syncytial 4-cell spermatheca-uterine valve (sp-ut valve). During ovulation, gonadal sheath cells contract and the distal neck of the spermatheca is pulled open to allow entry of the oocyte. After a regulated period of time the sp-ut valve opens and the distal spermathecal bag constricts to expel the embryo [11,12]. Failure to coordinate these distinct contraction events during ovulation leads to embryos that fail to reach the uterus, become misshapen, or have decreased viability [4,7].

The signaling pathways that regulate acto-myosin contractility in the spermatheca are similar to those found in smooth muscle and non-muscle cells [13–15]. Two pathways, a Ca^2+^ dependent and a Rho-dependent pathway are both necessary for spermathecal contractility. These pathways culminate in the phosphorylation and activation of non-muscle myosin. The Ca^2+^ dependent pathway relies on the activation of PLC-1/phospholipase C-ε, which cleaves phosphatidyl inositol (PIP_2_) into the second messengers diacylglycerol (DAG) and inositol 1,4,5 triphosphate (IP_3_). IP_3_ stimulates the release of Ca^2+^ from the endoplasmic reticulum (ER) through the ITR-1/IP_3_ receptor [3,5]. Cytosolic Ca^2+^ activates MLCK-1/myosin light chain kinase, which phosphorylates and activates myosin [16]. The Rho-dependent pathway is activated by oocyte entry, which displaces a mechanosensitive Rho GAP, SPV-1, from the membrane leading to the activation of RHO-1 [7]. GTP-bound RHO-1 then activates LET-502/ROCK, which in turn phosphorylates myosin and inhibits the myosin phosphatase subsequently increasing the levels of phosphorylated myosin [5,7]. The sp-ut valve opens as the bag constricts to push the embryo into the uterus. Although Ca^2+^ signaling and actin-based contractility are critical [4,5,16–20], little is known about the mechanisms regulating the timing and spatial coordination of Ca^2+^ release during embryo transit through the spermatheca.

In order to better understand the regulation of Ca^2+^ signaling in the spermatheca, we performed a candidate RNA interference (RNAi) screen and identified the heterotrimeric G-protein alpha subunit GSA-1/Gα_s_ as an important regulator of spermathecal Ca^2+^ release. Gα_s_ is a GTPase that regulates diverse biochemical responses in multiple tissue types, including the contractility of airway and arteriole smooth muscle cells [21,22]. In *C. elegans*, GSA-1 regulates an array of behaviors including larval viability, egg laying, oocyte maturation and locomotion [23–26]. GSA-1 is expressed in neurons, muscle cells, pharynx and the male tail [26–28] and throughout the somatic gonad, including the sheath cells, the spermatheca, uterus and the vulval muscles [24,26], however, the role of GSA-1 in the spermatheca has not been studied.

Heterotrimeric G proteins are composed of α, β, and γ subunits and are typically found downstream of the 7-pass transmembrane class of G-protein coupled receptors (GPCR). Upon ligand binding, the GPCR acts as a guanine nucleotide exchange factor (GEF), exchanging GDP for GTP on the α subunit, and activating the heterotrimeric G protein. Heterotrimeric G proteins can also be activated via a receptor-independent mechanism facilitated by G protein regulator proteins (GPRs) [21,29]. Upon activation, the GTP-bound α subunit disassociates from the βγ subunit and both the α and βγ subunits can initiate downstream signaling pathways [30]. *C. elegans* express two Gβ subunits, encoded by GPB-1 and GPB-2. GPB-1 shares 86% homology with mammalian β subunits and interacts with all Gα subunits in *C. elegans* [31,32]. There are two γ subunits, GPC-1 and GPC-2, in the *C. elegans* genome. GPC-1/γ is expressed in sensory neurons, while GPC-2/γ is expressed more broadly [33].Together, Gβγ subunit can regulate ion channels [34,35] including Ca^2+^ channels [36], as well as activate or inhibit adenylyl cyclase [37]. GPB-2 acts downstream of the Gα subunit GOA-1 in pharyngeal pumping [38], and is required for egg-laying and locomotion [39,40].

In smooth muscle, activation of Gα_s_ inhibits acto-myosin contractility through the activation of adenylyl cyclase, which converts adenosine triphosphate (ATP) to 3’-5’-cyclic adenosine monophosphate (cAMP). cAMP levels are in part regulated by phosphodiesterases (PDEs), which convert cAMP into AMP. PKA is composed of two catalytic (PKA-C) and two regulatory (PKA-R) subunits, and is activated when cAMP binds to the PKA-R subunits, causing the release of PKA-C. Therefore, PKA-R acts as an inhibitor of PKA-C in the absence of cAMP. Once released, PKA-C interacts with multiple downstream effectors and regulates lipid metabolism [41], cell migration [42], and vasodilation [43], among many other functions [44]. In airway smooth muscle, PKA inhibits contraction by antagonizing Gα_q_ mediated increases in Ca^2+^, as well as by phosphorylating and inhibiting Rho [45] and other regulators of contractility. In *C. elegans*, PKA is encoded by catalytic subunit *kin-1/*PKA-C and regulatory subunit *kin-2/*PKA-R. KIN-1 has been implicated in a variety of functions, such as cold tolerance [46], oocyte meiotic maturation [47], locomotion [48], immune responses [49], and Ca^2+^ influx into motor neurons [50].

Here we describe a role for GSA-1/Gα_s_ and its downstream effectors, KIN-1/PKA-C and KIN-2/PKA-R, in the regulation of Ca^2+^ release and contractility in the *C. elegans* spermatheca. Depletion of these proteins results in abnormal Ca^2+^ signaling and loss of coordinated contractility between the sp-ut valve and the spermathecal bag. Additionally, we describe a role for Gβ subunits GPB-1 and GPB-2 and phosphodiesterase PDE-6 in regulating spermathecal contractility and Ca^2+^ signaling.

## Results

### GSA-1/Gα_s_ and KIN-1/PKA-C are necessary for proper oocyte transit through the spermatheca

To determine if GSA-1/Gα_s_ is required for oocyte transit, we depleted it using RNA interference (RNAi) in animals expressing GCaMP3 (GFP-calmodulin-M13 Peptide, version 3), a genetically encoded Ca^2+^ sensor, in the spermatheca [5,51]. Expression of GCaMP3 in the spermatheca provided both a visible GFP outline for scoring and a sensor for Ca^2+^ levels. The percentage of spermathecae occupied by an embryo was scored in adult animals. Depletion of factors that play a role in embryo transit through the spermatheca result in an increase in the percentage of occupied spermathecae compared to empty vector, or control RNAi, treated animals (18% occupied, n=300). We used *plc-1*/PLC-ε, a gene essential for embryo transit, as a positive control (100% occupied, n=214) [3–5] (Fig. 1A).

**Figure 1:**
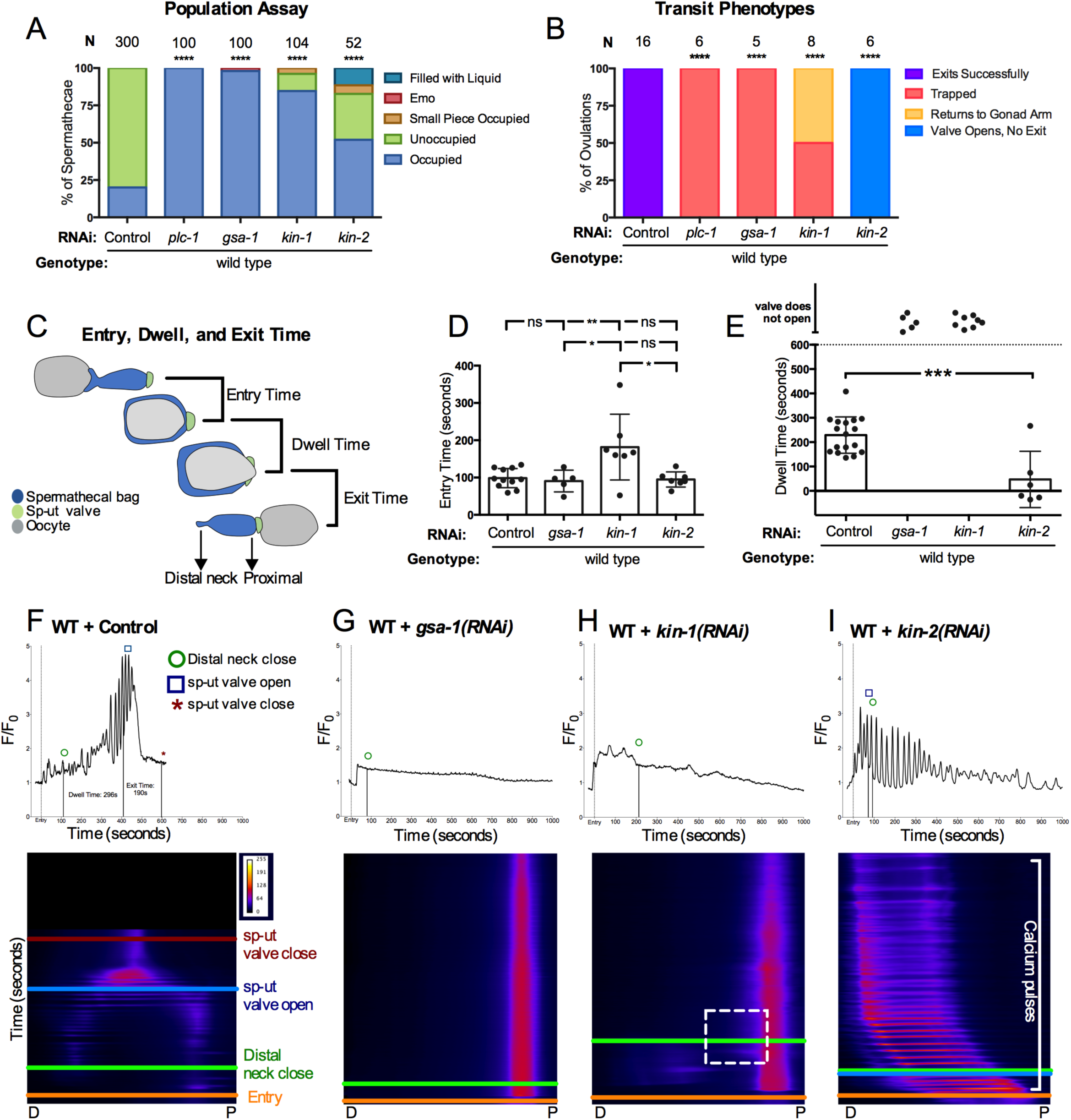
GSA-1/Gαs and KIN-1/PKA-C, and KIN-2/PKA-R regulate spermathecal contractility and Ca^2+^ dynamics in the spermatheca. (A) Population assay of wild type animals grown on control RNAi, *plc-1(RNAi), gsa-1(RNAi), kin-1(RNAi)*, and *kin-2(RNAi).* Spermathecae were scored for the presence or absence of an embryo (occupied or unoccupied), presence of a fragment of an embryo or presence of liquid (small piece occupied, filled with liquid), or the presence of endomitotic oocytes in the gonad arm (emo). The total number of unoccupied spermatheca were compared to the sum of all other phenotypes using the Fisher’s exact test. N is the total number of spermathecae counted. (B) Transit phenotypes of the ovulations of wild type animals treated with control RNAi, *gsa-1(RNAi), kin-1(RNAi)*, and *kin-2(RNAi)* and scored for successful embryo transits through the spermatheca (exits successfully), failure to exit (trapped), reflux into the gonad arm (returns to gonad arm), and the situation in which the sp-ut valve opens, but the embryo does not exit (valve opens, no exit). For transit phenotype analysis, statistics were performed using the total number of oocytes that exited the spermatheca successfully compared to the sum of all other phenotypes. Fisher’s exact tests were used for both population assays and transit phenotype analysis, and control RNAi was compared between all other RNAi treatments. (C) Schematic representation of a spermatheca undergoing an ovulation, with entry, dwell, and exit times indicated. Entry time (D) and dwell time (E) analysis of animals treated with control, *gsa-1(RNAi), kin-1(RNAi)*, and *kin-2(RNAi).* One-way ANOVA with a multiple comparison Tukey’s test was conducted to compare dwell times, and Fishers exact *t*-test (two dimensional *x*^*2*^ analysis) was performed on the exit times. Stars designate statistical significance (**** p<0.0001, *** p<0.005, ** p<0.01, * p<0.05). Representative normalized Ca^2+^ traces and kymograms of movies in B (F-G) are shown with time of entry, distal neck closure, and time the sp-ut valve opens and closes indicated. Levels of Ca^2+^ signal were normalized to 30 frames before oocyte entry. Kymograms generated by averaging over the columns of each movie frame (see methods).

Depletion of *gsa-1* by RNAi in GCaMP3 expressing animals resulted in almost complete spermathecal occupancy (98% occupied, n=61) with a low penetrance entry defect, in which gonad arms were filled with endomitotically duplicating oocytes (EMO) [52] (2% EMO, n=61;Fig. 1A). The EMO phenotype occurs when mature oocytes fail to enter the spermatheca and re-enter the mitotic cycle, accumulating DNA [52]. Depletion of GSA-1 resulted in 90% spermathecal occupancy (Fig. 1B), with the other 10% exhibiting entry defects (n=87). The null allele *gsa-1(pk75)* results in larval arrest, and therefore cannot be used to study the role of GSA-1 in the spermatheca [25]. These results reveal a novel role for GSA-1 in the regulation of spermathecal contractility.

Because GSA-1/Gα_s_ often stimulates the production of cAMP and activation of PKA, we next explored a possible role for PKA in regulation of spermathecal contractility. KIN-1/PKA-C, is expressed in the spermathecal bag and the sp-ut valve [41]. Depletion of KIN-1/PKA-C resulted in significantly increased spermathecal occupancy (85% occupied, n=104; Fig. 1A). Depletion of the regulatory subunit KIN-2/PKA-R also resulted in increased spermathecal occupancy (52% occupied, n=52; Fig. 1A,). These results suggest that regulation of PKA activity is critical for successful transit of fertilized embryos through the spermatheca.

### GSA-1/Gα_s_ KIN-1/PKA-C, and KIN-2/PKA-R regulate spermathecal contractility and Ca^2+^ dynamics

To better understand how GSA-1/Gα_s_, KIN-1/PKA-C, and KIN-2/PKA-R regulate transit of oocytes through the spermatheca, we scored time-lapse movies of animals expressing GCaMP3 in the spermatheca for the presence of four transit phenotypes: successful exit of the embryo (Exited Successfully), retention of the embryo in the spermatheca (Trapped), reflux of the embryo into the oviduct (Returned to Gonad Arm), and failure of the embryo to exit, despite an open sp-ut valve (Valve Opens, No Exit). As expected, all control RNAi embryos exited the spermatheca (100% Exited Successfully, n=16; Fig. 1B,) and all *gsa-1(RNAi)* embryos failed to exit the spermatheca (100% Trapped, n=5). Unlike wild type ovulations (Supplemental Movie 1), in *gsa-1(RNAi)* animals, the embryo remained in the spermatheca with the distal neck and sp-ut valve fully closed (Supplemental Movie 2). In *kin-1(RNAi)* animals, 50% of the embryos remained in the spermatheca, while the others returned to the gonad arm (n=8). We observed embryos that entered, returned to the gonad arm, and re-entered the spermatheca multiple times during imaging, while the sp-ut valve remained closed [Supplemental Movie 3]. In 100% of the *kin-2(RNAi)* ovulations, the sp-ut valve opened, often prematurely, but the spermatheca failed to produce the contractions necessary for the embryo to be pushed into the uterus (n=6).

To further quantify the transit dynamics, we measured the entry time, dwell time, and exit time [16,53] in the time lapse movies. The entry time is the time from the beginning of oocyte entry to when the distal neck closes, fully enclosing the oocyte in the spermatheca. The dwell time is measured from time of distal neck closure to the point the sp-ut valve begins to open. The exit time is measured from sp-ut valve opening, through expulsion of the fertilized embryo into the uterus and full closure of the sp-ut valve (Fig. 1C). Entry times in *gsa-1(RNAi)* and *kin-2(RNAi)* did not differ significantly from wild type. However, because of the defects in distal neck closure discussed above, *kin-1(RNAi)* animals had a significantly increased entry time (Fig. 1D). Because the sp-ut valve never opens in *gsa-1(RNAi)* and *kin-1(RNAi)* animals, neither dwell time nor exit time could be measured. In contrast, in *kin-2(RNAi)* animals, the sp-ut valve opens significantly faster than WT, resulting in low, and sometimes negative, dwell times (Fig. 1E). Despite these short dwell times, the oocytes consistently failed to exit the spermatheca during the 30-minute imaging period, resulting in no exit times for *kin-2(RNAi)* animals.

Taken together, these results are consistent with the hypothesis that PKA negatively regulates contractility in the spermatheca and sp-ut valve. We propose that when KIN-1/PKA-C is depleted, increased contractility in the proximal bag and sp-ut valve prevent exit of the fertilized embryo. In this case, the embryo is trapped or pushed back into the oviduct. Conversely, when KIN-2/PKA-R is depleted, activating KIN-1/PKA-C, neither the spermathecal bag nor sp-ut valve is contractile, resulting in reduced exit of the fertilized embryos. In this case, the embryo is not pushed out.

Because Ca^2+^ signaling plays a key role in spermatheca contractility [5], we next explored the role of GSA-1 and PKA in spermathecal Ca^2+^ dynamics. We collected time-lapse fluorescent images of animals expressing GCaMP3 and plotted the data both as 1D traces of total Ca^2+^ levels, and as 2D kymographs with time on the y-axis and space in the x-axis (see methods), which allows visualization of the spatial and temporal aspects of Ca^2+^ signaling. In wild type animals, the sp-ut valve exhibits a bright pulse of Ca^2+^ immediately upon oocyte entry, which is followed by a quiet period. Once the oocyte is completely enclosed, Ca^2+^ oscillates across the spermatheca, increasing in intensity until peaking concomitantly with distal spermathecal constriction and embryo exit [5,53] (Supplemental Movie 1, Fig. 1F). Depletion of *gsa-1(RNAi)* resulted in low levels of Ca^2+^ fluorescence in the spermathecal bag, which failed to increase over baseline levels. In contrast, Ca^2+^ signal in the sp-ut valve remained high throughout the entire observation period (n=5; Fig. 1G; Supplemental Movie 2). This suggests GSA-1 is required to initiate Ca^2+^ signaling in the spermathecal bag and for the correct inhibition of Ca^2+^ in the sp-ut valve.

Consistent with the phenotype observed in *gsa-1(RNAi)* animals, the Ca^2+^ signal in the distal spermathecal bag of *kin-1(RNAi)* animals failed to increase over baseline levels. However, some Ca^2+^ signal was observed on the proximal side of the spermathecal bag (n=8; Fig. 1H, indicated by box; Supplemental Movie 2). This increased proximal Ca^2+^ signal (Fig. 1H) coincided with retrograde motion of the embryo back into the gonad arm (see Fig. 1B; Supplemental Movie 3).

We next explored the effect of hyperactivation of the catalytic subunit KIN-1/PKA-C by depleting the regulatory subunit, KIN-2/PKA-R. Because depletion of KIN-1 results in a loss of Ca^2+^ signaling in the spermatheca bag, we expected that the loss of the regulatory subunit, KIN-2/PKA-R, might increase Ca^2+^ signaling by allowing the catalytic subunit to remain uninhibited. In *kin-2(RNAi)* animals, Ca^2+^ signaling increased immediately upon oocyte entry (Fig. 1I; Supplemental Movie 4). Ca^2+^ transients propagated from the distal to the proximal side of the spermatheca in rhythmic succession. These Ca^2+^ transients appear as pulses in the Ca^2+^ trace data and as horizontal lines in the kymogram (Fig. 1I). Although *kin-2(RNAi)* stimulated Ca^2+^ release, embryos failed to exit the spermatheca (n=7, Fig. 1B). These results suggest unregulated activity of KIN-1/PKA-C, through loss of KIN-2/PKA-R, results in abnormal Ca^2+^ signaling in the spermatheca, which is insufficient to stimulate embryo exit.

### GSA-1(GF) induction of Ca^2+^ signaling in the spermatheca is dependent on KIN-1/PKA-C and PLC-1

We next explored the effects of activating GSA-1 signaling on spermathecal transits. The allele *gsa-1(ce94)* is a gain of function allele (referred to here as GSA-1(GF)) in which the GTP-bound form of GSA-1 is stabilized, resulting in an elevated level of GSA-1 activity [54]. First, we analyzed transit parameters in the GSA-1(GF) animals. Most fertilized embryos in GSA-1(GF) animals passed successfully through the spermatheca, however, in 31% of transits, the sp-ut valve opened, but the embryo was unable to exit (n=13; Fig. 2A). Although entry times and dwell times in GSA-1(GF) animals did not differ from wild type (Fig. 2B), the exit times of GSA-1(GF) ovulations were of significantly longer duration, suggesting that the bag may not contract with the timing or force needed to efficiently expel the embryo into the uterus (Fig. 2C). We reasoned that if PKA is downstream of GSA-1, depletion of KIN-1/PKA-C in the GSA-1(GF) background would result in increased trapping of embryos in the spermatheca. Indeed, GSA-1(GF) animals treated with *kin-1(RNAi)* resulted in a complete failure of embryos to exit the spermatheca. We observed 67% of the embryos remained entirely enclosed by the spermatheca, while in the remaining 33% the valve opened but the embryo was unable to exit (n=6; Fig. 2A).

**Figure 2:**
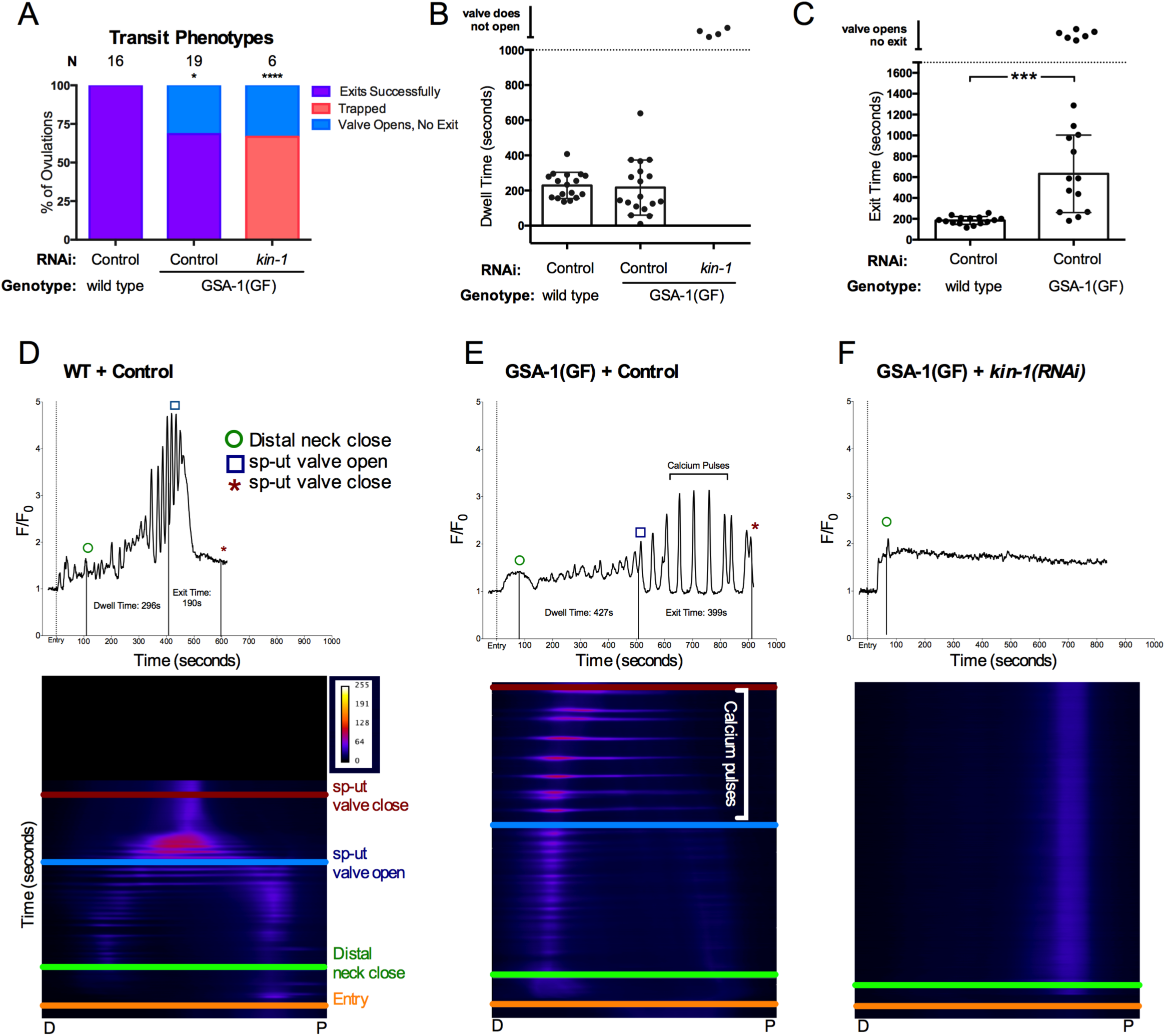
GSA-1(GF) induction of Ca^2+^ signaling in the spermatheca is dependent on KIN-1/PKA-C. (A) Transit phenotypes of the ovulations of wild type animals treated with control RNAi, and GSA-1(GF) animals treated with control RNAi and *kin-1(RNAi)*, were scored for successful embryo transits through the spermatheca (exits successfully), failure to exit (trapped), reflux into the gonad arm (returns to gonad arm), and the situation in which the sp-ut valve opens, but the embryo does not exit (valve opens, no exit). The total number of oocytes that exited the spermatheca successfully was compared to the sum of all other phenotypes using the Fisher’s exact test. Control RNAi was compared to all other RNAi treatments. Dwell (B) and exit (C) times of movies in A were analyzed using Fishers exact *t*-test (two dimensional *x*^*2*^ analysis). Stars designate statistical significance (**** p<0.0001, *** p<0.005, ** p<0.01, * p<0.05). Representative normalized Ca^2+^ traces and kymograms of movies in A (D-F) are shown with time of entry, distal neck closure, and time the sp-ut valve opens and closes indicated. Levels of Ca^2+^ signal were normalized to 30 frames before oocyte entry. Kymograms generated by averaging over the columns of each movie frame (see methods).

Because *gsa-1(RNAi)* reduced Ca^2+^ signaling in the spermathecal bag but increased signaling in the sp-ut valve, we predicted increasing GSA-1 activity would produce the opposite effect. To study the effect of GSA-1(GF) on ovulation and spermathecal Ca^2+^ dynamics, we collected time-lapse ovulation movies of GCaMP3-expressing GSA-1(GF) animals. As previously discussed, during WT ovulations, Ca^2+^ gradually increases, peaking concomitantly with spermathecal exit (Fig. 2D). However, rather than exhibiting prematurely elevated Ca^2+^, expression of GSA-1(GF) resulted in delayed Ca^2+^ release in the bag (n=16; Fig. 2E). The signal remained only slightly over baseline for the first few hundred seconds, after which Ca^2+^ was released in pulses that continued until embryo exit. In the sp-ut valve, Ca^2+^ signal was depressed, as expected (Compare Fig. 2D and 2E). The Ca^2+^ pulses in the bag sweep distally to proximally across the spermathecal tissue, and resemble the Ca^2+^ dynamics observed when PKA activity is increased through the depletion of KIN-2/PKA-R (Compare Fig. 1I to Fig. 2E). The similarity between *kin-2(RNAi)* animals and GSA-1(GF) animals suggests GSA-1 is acting through PKA in the spermatheca. Therefore, we predicted that depletion of KIN-1/PKA-C would suppress the Ca^2+^ phenotypes seen in the GSA-1(GF) animals. Indeed, depletion of KIN-1 in GSA-1(GF) prevented the Ca^2+^ pulses observed in GSA-1(GF) animals, resulting in a flat signal (Fig. 2F). The sp-ut valve failed to open, and the Ca^2+^ traces resembled those seen in *kin-1(RNAi)* animals. These results suggest KIN-1 is acting downstream of GSA-1 in the spermatheca to regulate Ca^2+^ release.

In wild type animals, oocyte entry stimulates Ca^2+^ release [5]. The spermathecal Ca^2+^ remains at baseline until ovulation, and returns to baseline following embryo exit (Fig. 3A). However, while observing GSA-1(GF) animals, we noticed dynamic changes in Ca^2+^ in unoccupied spermathecae (Fig. 3B and Fig. 3C). Similarly, in *kin-2(RNAi)* animals, Ca^2+^ repeatedly increases, peaks, and then drops to baseline levels in empty spermathecae (Fig. 3D). These pulses travel from the distal spermatheca through the bag to the sp-ut valve (Fig. 3C, D). These results suggest that activating GSA-1 or PKA can bypass the need for oocyte entry as a trigger for Ca^2+^ release. The similarities between GSA-1(GF) and *kin-2(RNAi)* further support the hypothesis that GSA-1 triggers Ca^2+^ signaling through KIN-1/KIN-2.

**Figure 3:**
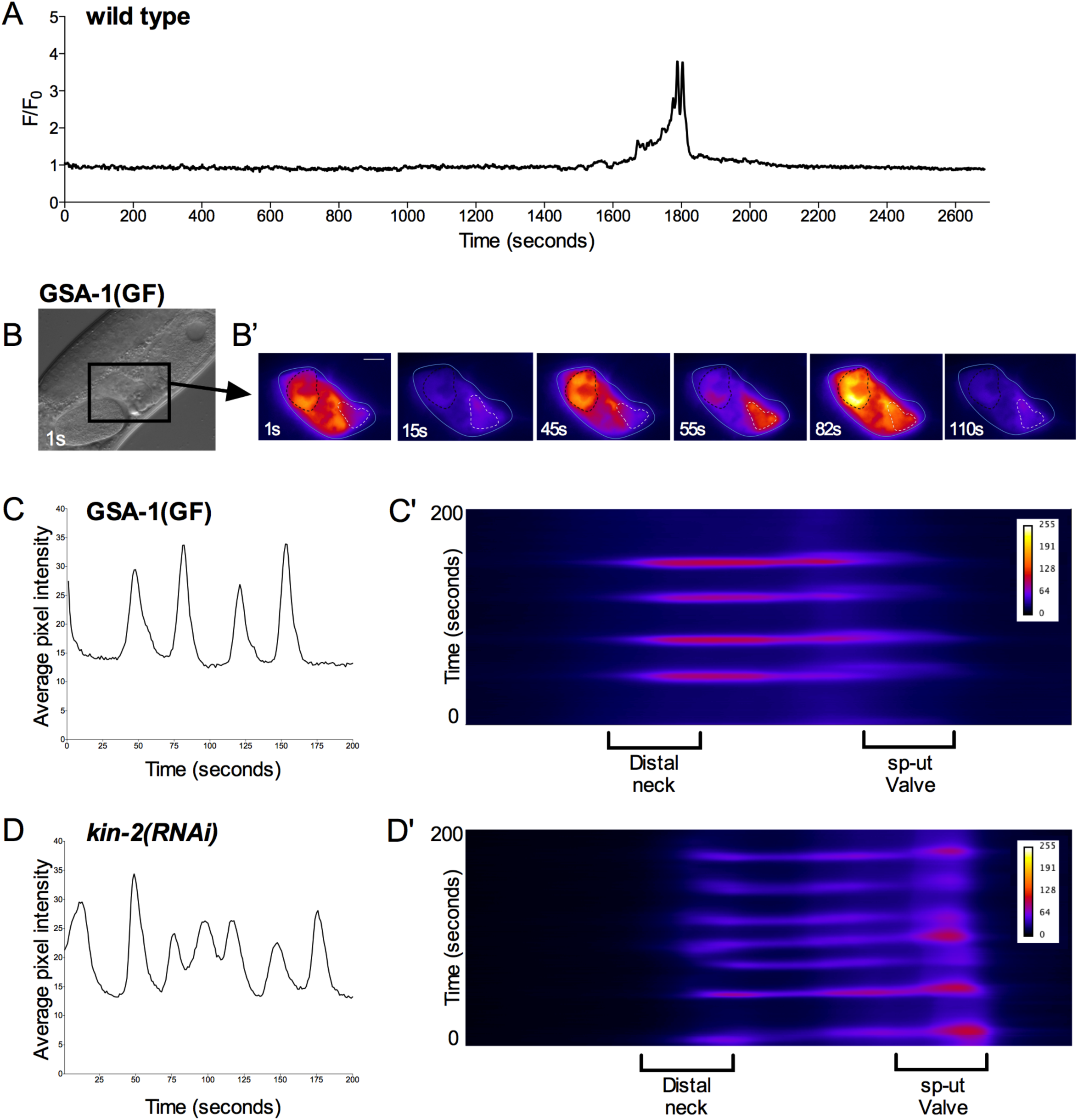
GSA-1(GF) and knockdown of KIN-2/PKA-R induces Ca^2+^ signaling in the empty spermatheca. (A) Normalized Ca^2+^ trace of a wild type ovulation before, during, and after transit. Ca^2+^ signal does not rise above basal levels before or after transits. (B) DIC image of a GSA-1(GF) animal; the spermatheca is indicated by a box. (B’) Ca^2+^ signaling in the unoccupied GSA-1(GF) spermatheca shown in B. Ca^2+^ repeatedly increases, peaks, and then drops to baseline levels. These pulses travel from the distal spermatheca through the bag to the sp-ut valve. Average pixel intensity trace of the Ca^2+^ signal (C) and kymogram (C’) of the GSA-1(GF) spermatheca are shown. Knockdown of KIN-2 with RNAi results in a similarly pulsing unoccupied spermatheca, as shown by the average pixel intensity trace of the Ca^2+^ signal (D) and kymogram (D’) of a *kin-2(RNAi)* animal.

The phospholipase PLC-1 is required for spermathecal contractility and Ca^2+^ release in the spermatheca [5]. Therefore, we next asked whether the phospholipase PLC-1 was required for transits in GSA-1(GF) animals. As expected, the null allele *plc-1(rx1)* resulted in 100% spermathecal occupancy (n=7; Fig. 4A). Similarly, in GSA-1(GF) animals treated with *plc-1(RNAi)*, embryos did not exit the spermatheca, but the sp-ut valve did open (n=4; Fig. 4A). Occasionally, the valve opened before the distal neck closed, resulting in negative dwell time (Fig. 4B). We next asked if the Ca^2+^ signal observed in the GSA-1(GF) animals (Fig. 4C) required phospholipase-stimulated Ca^2+^ release. In *plc-1* null animals, Ca^2+^ signaling remains at baseline levels in the spermathecal bag even after oocyte entry (Fig.6D). When GSA-1(GF) animals were treated with *plc-1(RNAi*), the Ca^2+^ signal remained low in the spermathecal bag, similar to the levels observed in *plc-1(rx1)* animals, and we no longer observed the Ca^2+^ pulses characteristic of GSA-1(GF) (Fig. 4E). Similarly, loss of *plc-1* prevented both the excess proximal Ca^2+^ release observed in *kin-1(RNAi)* spermathecae (Fig. 4F), and the Ca^2+^ pulses observed in *kin-2(RNAi)* spermathecae (Fig. 4G). These results suggest the Ca^2+^ pulses observed when either GSA-1 or PKA signaling is activated require PLC-1-stimulated Ca^2+^ release, and indicate GSA-1 and PKA act upstream or in parallel to PLC-1 to stimulate Ca^2+^ release in the spermatheca.

**Figure 4:**
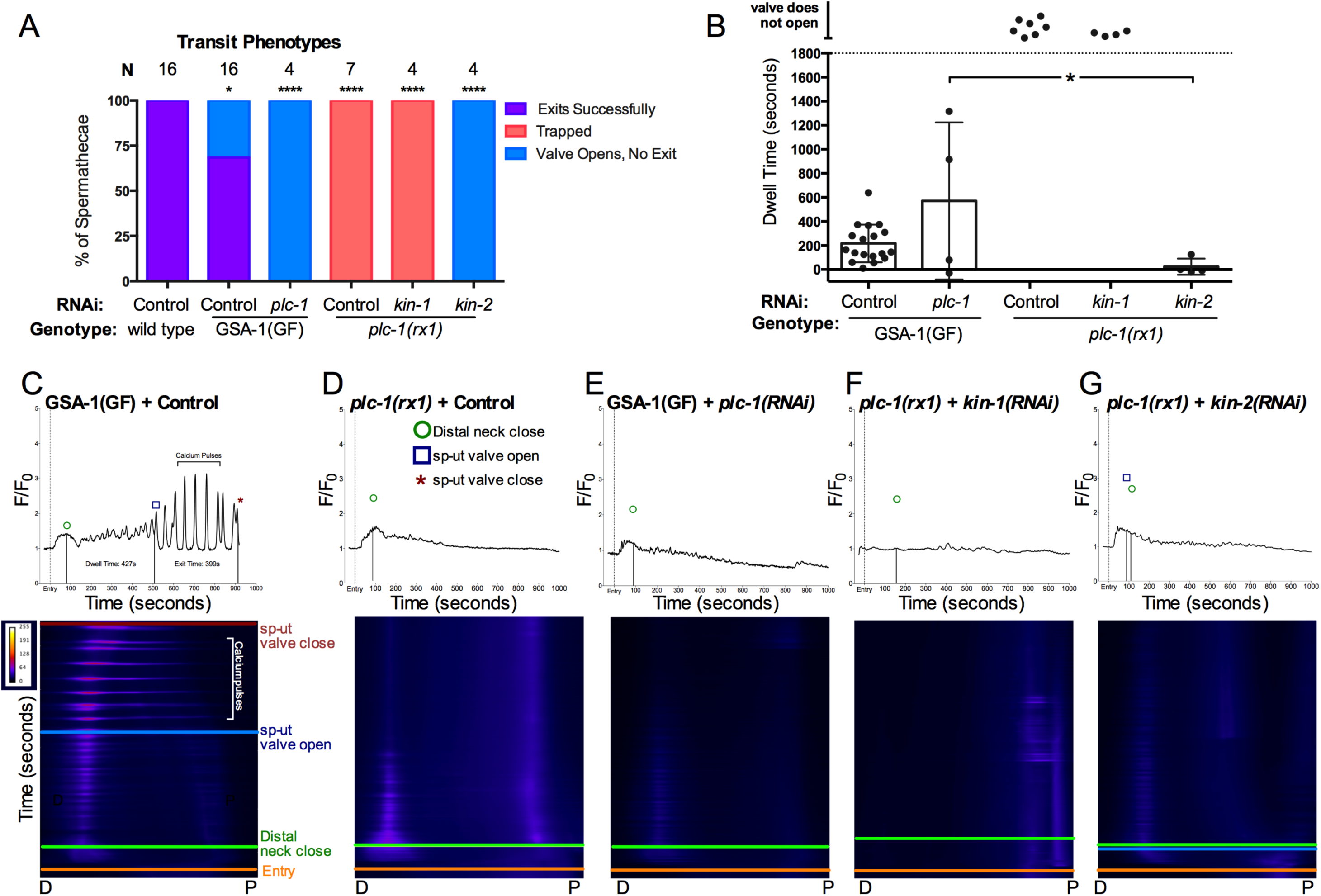
Ca^2+^ signal is dependent on PLC-1. (A) Transit phenotypes of the ovulations of wild type animals treated with control RNAi, GSA-1(GF) animals treated with control RNAi and *plc-1(RNAi)*, and *plc-1* null animals treated with control RNAi, *kin-1(RNAi), and kin-2(RNAi)* were scored for successful embryo transits through the spermatheca (exits successfully), failure to exit (trapped), and the situation in which the sp-ut valve opens, but the embryo does not exit (valve opens, no exit). The total number of oocytes that exited the spermatheca successfully was compared to the sum of all other phenotypes using the Fisher’s exact test. Control RNAi was compared to all other RNAi treatments. (B) Dwell times derived from movies in A were compared using One-way ANOVA with a multiple comparison Tukey’s test. Stars designate statistical significance (**** p<0.0001, *** p<0.005, ** p<0.01, * p<0.05). Representative normalized Ca^2+^ traces and kymograms of movies in A (C-G) are shown with time of entry, distal neck closure, and time the sp-ut valve opens and closes indicated. Levels of Ca^2+^ signal were normalized to 30 frames before oocyte entry. Kymograms generated by averaging over the columns of each movie frame (see methods).

Phosphodiesterase PDE-6 is necessary for normal transit times and Ca^2+^ dynamics Because PKA is activated by cAMP, we anticipated that altering cAMP levels in the spermatheca would alter the contractility and Ca^2+^ dynamics during ovulation. Phosphodiesterases (PDEs) are enzymes that regulate the concentration of cAMP by catalyzing its hydrolysis into AMP. To determine whether increasing cAMP levels by depleting PDEs would affect spermathecal transits, we depleted each of the known *C. elegans* PDEs and identified PDE-6 as an important regulator of spermathecal contractility. Depletion of *pde-6* resulted in 56% (n=50) of animals with embryos retained in the spermatheca (Fig. 5A). To study the effects of PDE-6 on ovulations and Ca^2+^ dynamics, we collected time-lapse images of GCaMP3 animals fed *pde-6(RNAi)*. Both the entry time and the exit time in *pde-6(RNAi)* animals were significantly longer than WT (Fig. 5B and Fig. 5C. In one case, the sp-ut valve never opened, and the embryo remained trapped in the spermatheca (Fig. 5D). Like GSA-1(GF) ovulations, *pde-6(RNAi)* ovulations resulted in delayed Ca^2+^ release in the bag, after which the Ca^2+^ was released in pulses until embryo exit (n=8; Fig. 5E). The Ca^2+^ pulses were reminiscent of those seen in both GSA-1(GF) and *kin-2(RNAi)*. These results suggest that a specific level of cAMP signaling through PKA regulates spermathecal contractility.

**Figure 5:**
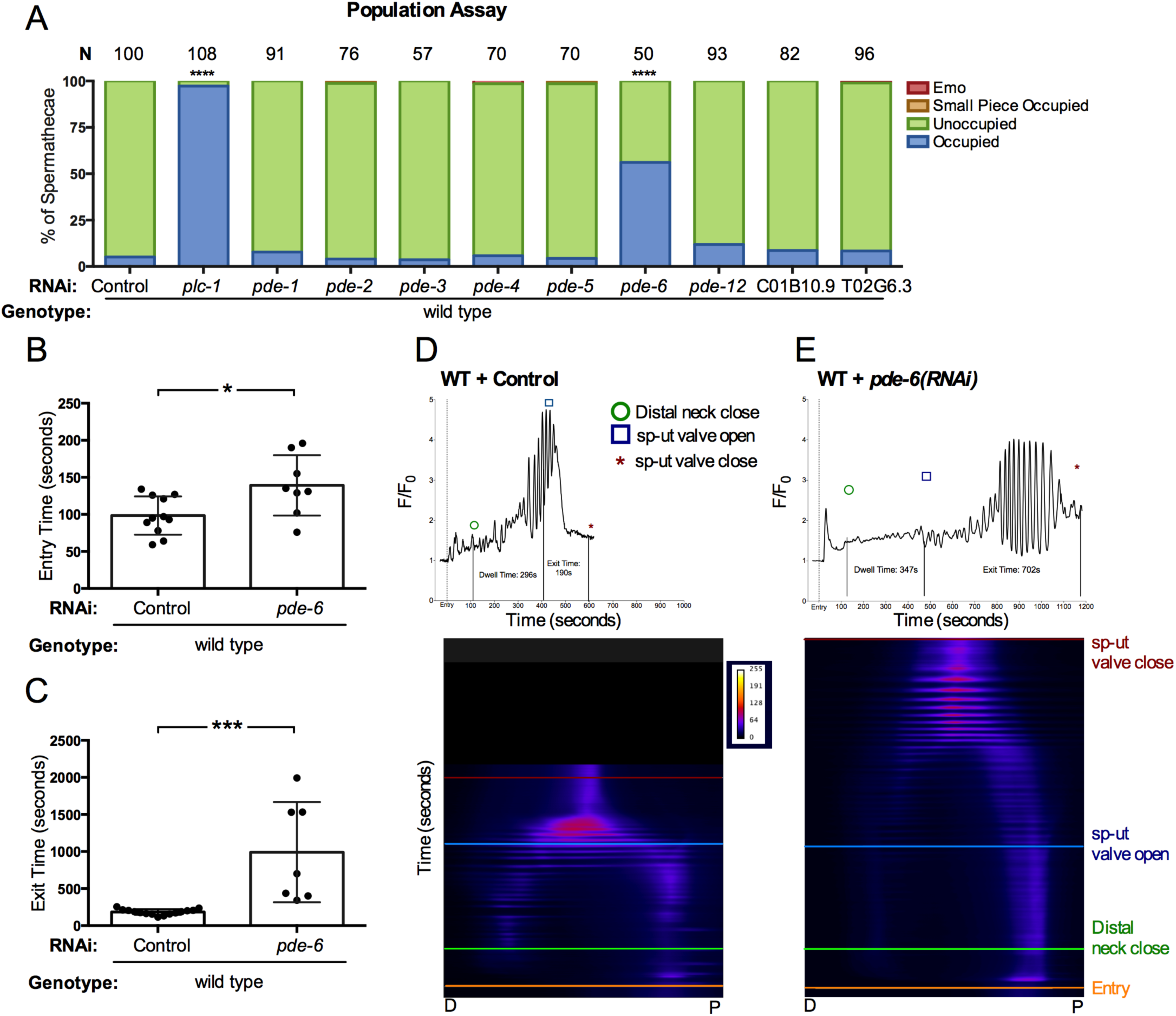
Phosphodiesterase PDE-6 is necessary for normal transit times and Ca^2+^ dynamics. (A) Population assay of wild type animals grown on control RNAi, *plc-1(RNAi), pde-1(RNAi), pde-2(RNAi), pde-3(RNAi), pde-4(RNAi), pde-5(RNAi), pde-6(RNAi), pde-12(RNAi)*, C01B10.9 and T02G6.3. Spermathecae were scored for the presence or absence of an embryo (occupied or unoccupied), presence of a fragment of an embryo (small piece occupied), or the presence of endomitotic oocytes in the gonad arm (emo) phenotypes. The total number of unoccupied spermatheca was compared to the sum of all other phenotypes using the Fisher’s exact test. N is the total number of spermathecae counted. Control RNAi was compared to all other RNAi treatments. Entry time (B) and exit time (C) were compared using Fishers exact *t*-test (two dimensional *x*^*2*^ analysis). Stars designate statistical significance (**** p<0.0001, *** p<0.005, ** p<0.01, * p<0.05). Representative normalized Ca^2+^ traces and kymograms of control RNAi (D) and *pde-6(RNAi)* (E) are shown with time of entry, distal neck closure, and time the sp-ut valve opens and closes indicated. Levels of Ca^2+^ signal were normalized to 30 frames before oocyte entry. Kymograms generated by averaging over the columns of each movie frame (see methods).

### Heterotrimeric G-protein beta subunit GPB-1 and GPB-2 are regulate Ca^2+^ signaling and spermatheca contractility

Heterotrimeric G proteins consist of an α, a β and a γ subunit. The β and γ subunits are closely associated and can considered one functional unit. Activation of the Gα subunit causes Gα to dissociate from Gβγ, and both can then initiate downstream signaling pathways [55]. Therefore, we next explored potential roles for βγ in the spermatheca. Depletion of GPB-1/β resulted in a significant increase in the percent of occupied spermathecae (34%, n=99) compared to control RNAi treated animals (18% occupied, n=300; Fig. 6A). GPB-1 depletion also resulted in a significant number of spermathecae occupied by a small piece of an embryo (Fig. 6A). Surprisingly, while GPB-2 depletion showed no significant trapping in freely moving animals (Fig. 6A), when the animals were immobilized for imaging, embryos failed to exit the spermatheca in 60% of *gpb-2(RNAi)* movies (n=10) (Fig. 6B), suggesting animals may be able to compensate somewhat for the loss of GPB-2 with body wall muscle contraction. Neither *gpb-1* nor *gpb-2* RNAi on their own affected the time required for the embryo to exit the spermatheca (Fig. 6C). Depletion of neither GPC-1/γ nor GPC-2/γ significantly affected ovulation when depleted via RNAi (Fig. 6A). Similarly, neither the *gpc-1(pk298)* null allele, nor treating *gpc-1(pk298)* with *gpc-2* RNAi resulted in ovulation defects. These data suggest GPB-1 may be associated with GSA-1 in the spermatheca (Fig. 6A), but identification of the relevant γ subunits requires further investigation. Insufficient knockdown of GPC-1/γ nor GPC-2/γ may explain the lack of phenotype.

**Figure 6:**
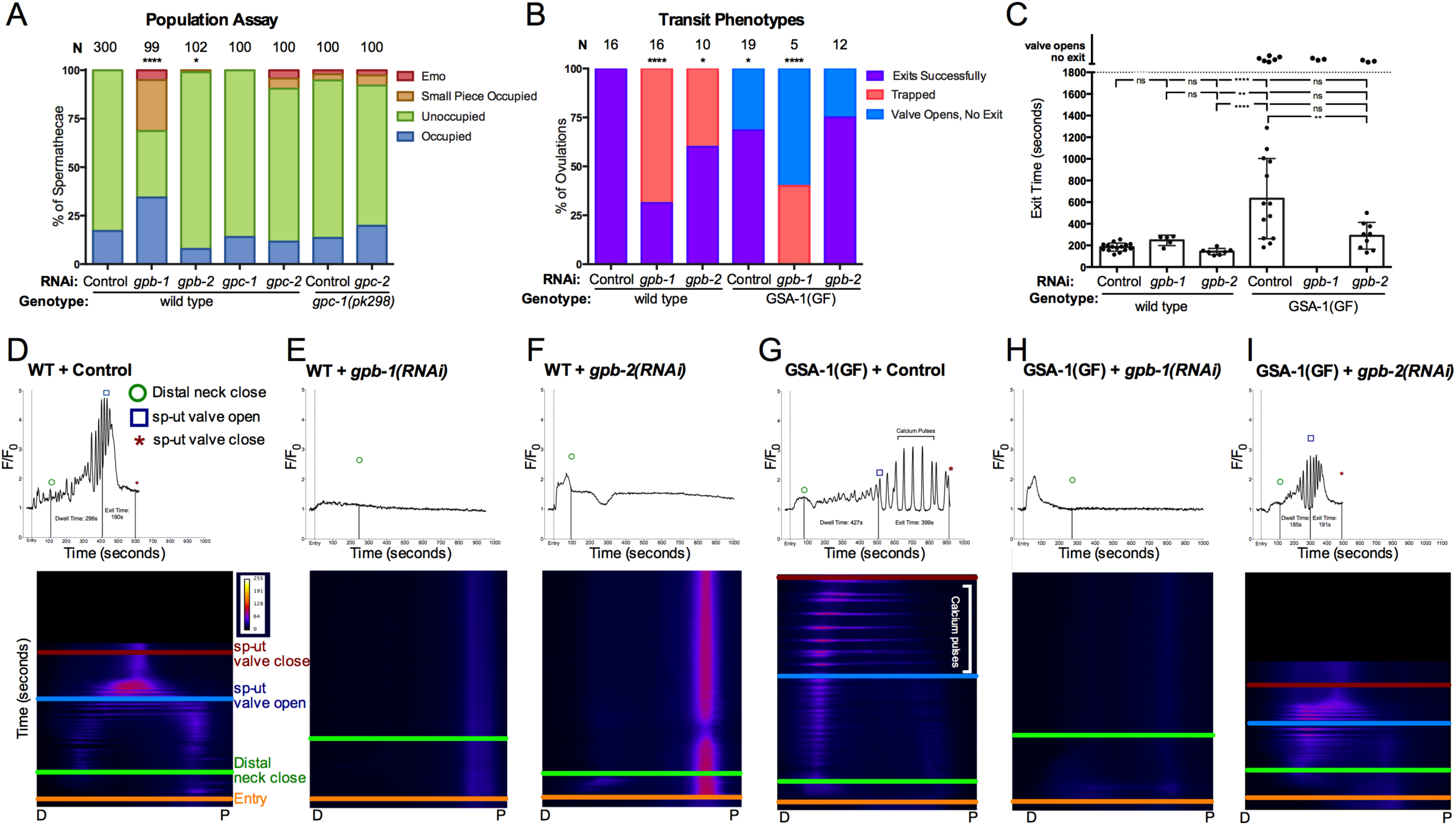
Heterotrimeric G-protein beta subunits GPB-1 and GPB-2 regulate Ca^2+^ signaling and spermathecal contractility. (A) Population assay of wild type young adults grown on control RNAi, *gpb-1(RNAi), gpb-2(RNAi), gpc-1(RNAi)*, and *gpc-2(RNAi)* and *gpc-1(pk298)* grown on control RNAi and *gpc-2(RNAi).* Spermathecae were scored for the presence or absence of an embryo (occupied or unoccupied), presence of a fragment of an embryo (small piece occupied), or the presence of endomitotic oocytes in the gonad arm (emo) phenotypes. The total number of unoccupied spermatheca was compared to the sum of all other phenotypes using the Fisher’s exact test. N is the total number of spermathecae counted. (B) Transit phenotypes of the ovulations in wild type animals treated with control RNAi, *gpb-1(RNAi)* and *gpb-2(RNAi)*, and GSA-1(GF) animals treated with control RNAi, *gpb-1(RNAi)*, and *gpb-2(RNAi)* were scored for successful embryo transits through the spermatheca (exits successfully), failure to exit (trapped), and the situation in which the sp-ut valve opens, but the embryo does not exit (valve opens, no exit). For transit phenotype analysis, the total number of oocytes that exited the spermatheca successfully was compared to the sum of all other phenotypes. Fisher’s exact tests were used for both population assays and transit phenotype analysis. (C) Exit time of movies in B were compared using One-way ANOVA with a multiple comparison Tukey’s test. Stars designate statistical significance (**** p<0.0001, *** p<0.005, ** p<0.01, * p<0.05). Representative normalized Ca^2+^ traces and kymograms of movies in B (D-I) are shown with time of entry, distal neck closure, and time the sp-ut valve opens and closes indicated. Levels of Ca^2+^ signal were normalized to 30 frames before oocyte entry. Kymograms generated by averaging over the columns of each movie frame (see methods).

In order to explore a role for GPB-1 and GPB-2 in Ca^2+^ signaling, we observed ovulations in GCaMP3 expressing animals treated with *gpb-1(RNAi)* and *gpb-2(RNAi)*. Compared to wild type movies, in which oocyte entry triggers a pulse of Ca^2+^ in the sp-ut valve, followed by Ca^2+^ oscillations in the spermathecal bag and sp-ut valve that increase in intensity until the embryo exits (Fig. 6D), both *gpb-1(RNAi)* and *gpb-2(RNAi)* resulted in elevated and/or prolonged Ca^2+^ signal in the sp-ut valve and very little Ca^2+^ signal in the spermathecal bag, accompanied by embryo trapping (Fig. 6E, Fig. 6F). These results phenocopy the Ca^2+^ phenotype observed in *gsa-1(RNAi)* and *kin-1(RNAi)*, suggesting GPB-1 and GPB-2 are required for the activation of Ca^2+^ signaling by GSA-1.

Activation of Gα also liberates the Gβγ subunits. In order to determine if activated signaling through Gβγ subunits could partially explain the GSA-1(GF) phenotypes, we depleted GPB-1 and GPB-2 in the GSA-1(GF) background. First, we assessed the effect on GSA-1(GF) transit phenotypes. In GSA-1(GF) animals, even though the sp-ut valve opens, ∼30% of embryos remain inside the spermatheca (n=19; Fig. 6B). Depletion of GPB-1 on its own results in significant trapping, with most spermathecae occupied with an embryo (36%) or an embryo fragment (26%) (n=99; see Fig. 6A). When GPB-1 is depleted in the GSA-1(GF) background, no successful transits are observed, with 40% of ovulations resulting in trapping, and 60% of ovulations in which the valve opened but the embryo was unable to exit (n=5; Fig. 6B). In contrast, in GSA-1(GF) *gpb-2(RNAi)* animals, the trapping phenotype does not differ significantly from GSA-1(GF) alone (Fig. 6B). However, the significantly longer exit time observed in GSA-1(GF) animals is suppressed by *gpb-2(RNAi)* to nearly WT levels (Fig. 6C).

As previously described, GSA-1(GF) results in strong Ca^2+^ pulses that propagate across the tissue (Fig. 6G). When GPB-1 was depleted in GSA-1(GF) expressing animals, other than a brief pulse upon entry, little signal was observed in either the sp-ut valve or in the spermatheca bag (Fig. 6H). The similarities between the *gpb-1(RNAi)* and *gsa-1(RNAi)* phenotypes in the spermathecal bag suggest that even the gain of function allele of GSA-1 requires GPB-1 for sufficient activity to drive Ca^2+^ signaling. These results are consistent with the model that the binding to the Gβγ subunit is required for activation of GSA-1. In contrast, *gpb-2(RNAi)* suppressed the GSA-1(GF) Ca^2+^ phenotype nearly to wild type, with low-level Ca^2+^ oscillations increasing in intensity until embryo exit (Fig. 6I). Because Gβγ subunits are needed for the activation of Gα, depleting Gβ would be expected to reduce Gα activity. In a Gα hyperactive background, this could bring Gα signaling to a more normal level, therefore explaining the rescued exit times and Ca^2+^ signaling. Alternatively, the Gβs could be activating parallel signaling pathways needed for Ca^2+^ release, potentially including other Gα subunits active in the spermatheca (Reference Fig. 7).

**Figure 7:**
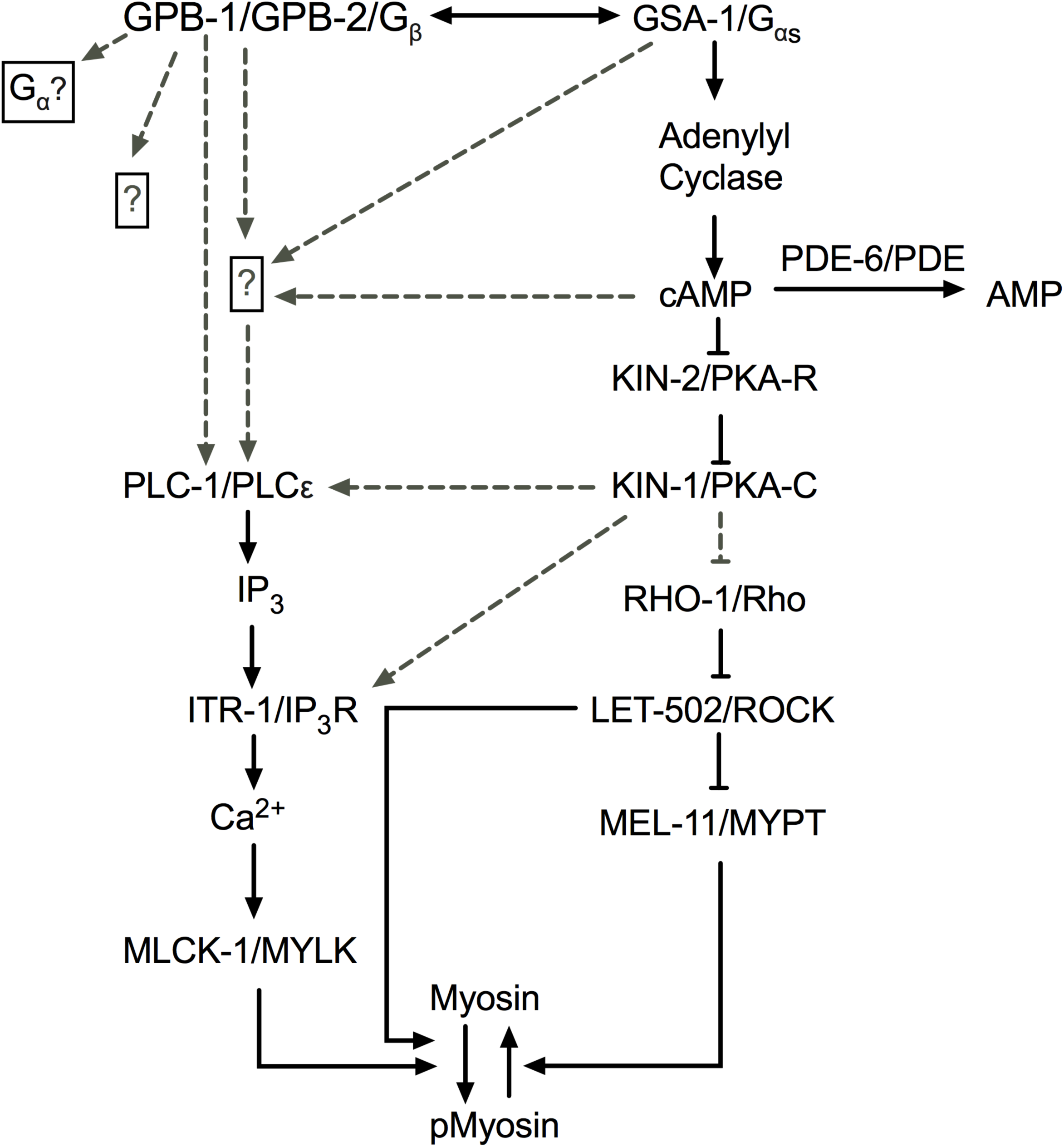
Proposed signaling network in the spermatheca. A working model of the spermathecal Ca^2+^ signaling network, with dashed lines indicating possible interactions. We propose KIN-1 and KIN-2 are working downstream of GSA-1, while GPB-1 and GPB-2 are either working together with or in parallel to GSA-1 to regulate Ca^2+^ signaling and spermathecal contractility.

## Discussion

Ca^2+^ and cAMP are important second messengers with key roles in a diverse set of biological processes. Here, we show that GSA-1/Gα_s_ and KIN-1/PKA-C function to regulate Ca^2+^ release and coordinated contractility in the *C. elegans* spermatheca. The G_β_ subunits GPB-1 and GPB-2 are required for activation of GSA-1/Gα_s_, and phosphodiesterase PDE-6 functions in the spermatheca to regulate Ca^2+^ levels and overall tissue contractility.

In the spermatheca, oocyte entry triggers Ca^2+^ waves, which propagate across the spermatheca and trigger spermathecal contractility. We found that loss of either GSA-1/Gα_s_ or KIN-1/PKA-C leads to low levels of Ca^2+^ signal in the spermathecal bag, while increasing GSA-1 activity with a gain of function allele or activating PKA through loss of the regulatory subunit, KIN-2/PKA-R, leads to strong Ca^2+^ pulses, even in the absence of oocyte entry. This suggests GSA-1/Gα_s_ and KIN-1/PKA-C stimulate Ca^2+^ release in the spermatheca. Previous work in our lab has shown that PLC-1/phospholipase C-ε is necessary to stimulate Ca^2+^ release in the spermatheca [5]. Here, we show that loss of PLC-1 can block the Ca^2+^ release observed in GSA-1(GF) and *kin-2(RNAi)* animals, suggesting PLC-1 acts either downstream or in parallel with Gα_s_ and PKA (Fig. 7). Mechanisms by which PKA could stimulate Ca^2+^ release include activation of the ITR-1/IP3 receptor, or through activation of plasma membrane channels such as stretch-sensitive TRPV channels [56]. For example, in mouse cardiomyocytes, Gα activation can stimulate Ca^2+^ release through exchange protein directly activated by cAMP (EPAC) and Rap1 [57,58]. However, we observed no phenotype when the *C. elegans* homologs of these factors were depleted by RNAi (data not shown), which may suggest a different mechanism of Ca^2+^ release in the spermatheca. In COS-7cells, PLC-ε can be activated by Gβγ subunits [59], which offers a promising alternative. Future work is needed to explore a possible connection between Gβγ and PLC-1 in the spermatheca.

Contraction of the spermatheca requires both Ca^2+^ and Rho signaling. Therefore, the observed transit defects could be due to GSA-1/Gα_s_ and KIN-1/PKA-C regulation of either, or both, of these signaling pathways. Because *kin-2(RNAi)*, which activates KIN-1/PKA-R and stimulates Ca^2+^ release, is unable to produce the correct contraction, this implies PKA may also regulate the Rho side of the pathway. In SH-EP cells, PKA has been shown to phosphorylate and inhibit Rho through stabilization of the inactive Rho GDI state [45,60] Given that the Rho phosphorylation site is conserved in the *C. elegans* RHO-1, it is possible that knockdown of KIN-2 results in an increased inhibition of RHO-1 thus resulting in decreased tissue contractility in *kin-2(RNAi)* animals.

The sp-ut valve is a syncytium of 4 cells that prevents premature release of oocytes into the uterus upon entry in the spermatheca. This allows for enough time for the oocyte to become fertilized and form an eggshell, after which the sp-ut dilates and allows passage of the embryo unto the uterus [61]. Here, we show that PKA regulates Ca^2+^ release and valve contractility in the sp-ut valve. Depletion of KIN-1/PKA-C activity results in increased Ca^2+^ and a valve that remains closed. In contrast, hyperactive PKA-C activity (through *kin-2(RNAi)*) results in an sp-ut valve that opens prematurely, perhaps accounting for the short dwell times observed when PKA-R is depleted (see Fig. 1E). Decreasing signal through PKA/KIN-1 or Gα_s_ /GSA-1 knockdown results in a surplus of Ca^2+^ in the sp-ut valve, whereas increasing signaling through GSA-1(GF) results in an extended quiet Ca^2+^ period in the sp-ut valve. This suggests PKA/KIN-1 and Gα_s_ /GSA-1 are required for Ca^2+^ inhibition in the sp-ut valve even though they are required for Ca^2+^ release in the spermathecal bag. Little is known about the signaling networks that regulate contractility in the sp-ut valve. PLC-1 is not expressed in the sp-ut valve [3], necessitating a different mechanism of Ca^2+^ regulation from the spermathecal bag. Phosphorylation of IP_3_Rs by PKA-C has been shown to decrease Ca^2+^ release in rat brain cells [62], which could help to explain the Ca^2+^ excess seen in the sp-ut valve. Other mechanisms by which PKA has been described to lower Ca^2+^ include increasing the activity of SERCA pumps, which pump Ca^2+^ back into the ER, by phosphorylating and dissociating phospholamban [63,64]. However, there is no obvious phospholamban homolog in *C. elegans*. PKA can inhibit PLC-β [65], which would result in decreased Ca^2+^ release but EGL-8/PLC-β is not expressed in either the spermatheca or sp-ut valve [66]. Future study may reveal the details of how PKA regulates Ca^2+^ signaling in the sp-ut valve.

Regulating the correct force, timing, and direction of contraction in biological tubes is crucial for living organisms. Dysfunction in the regulatory networks controlling these dynamics lead to diseases such as heart disease and asthma [43,67]. The signaling networks that control Ca^2+^ oscillations and actomyosin contractility in smooth muscle in humans are conserved in the *C. elegans* spermatheca. In this study, we identified Gα_s_ and PKA as key players in the spermatheca, where Gα_s_ acts upstream of PKA to regulate the spatiotemporal regulation of Ca^2+^ release and contractility in the spermatheca. Because PKA is implicated in a variety of biological processes, understanding its control and targets may give us insights into the diseases that occur when this control goes awry.

## Materials and Methods

### Strains and Culture

Nematodes were grown on nematode growth media (NGM) (0.107 M NaCl, 0.25% wt/vol Peptone (Fischer Science Education), 1.7% wt/vol BD Bacto-Agar (FisherScientific), 0.5% Nystatin (Sigma), 0.1 mM CaCl_2_, 0.1 mM MgSO_4_, 0.5% wt/vol cholesterol, 2.5 mM KPO_4_) and seeded with *E. coli OP50* using standard *C. elegans* techniques [68]. Nematodes were cultured at 23°C unless specified otherwise. All Ca^2+^ imaging was acquired using the strain UN1108 *xbls 1101 [fln-1p::GCaMP3,rol-6]* except for experiments utilizing the GSA-1 gain of function mutant KG524 *GSA-1(GF).* These animals were crossed into UN1417, a separate integration event of *xbIs 1108 [fln-1p::GCaMP3]*.

### Construction of the transcriptional and translational reporter of GSA-1

The *gsa-1* promoter (1.6 kb upstream of GSA-1 start codon) was amplified from *C. elegans* genomic DNA using primers with PstI and BamHI 5’ extensions and ligated into pPD95_77 (Fire Lab) upstream of GFP creating pUN783. To make a translational fusion *gsa-1* was amplified without its stop codon from *C. elegans* coding DNA using primers with BamHI 5’ extensions and ligated between the *gsa-1* promoter and GFP of pUN783 creating pUN810. Transgenic animals were created by microinjecting a DNA solution of 20 ng/μl of pUN783 or pUN810 and 50 ng/μl of pRF4 *rol-6* (injection marker) into N2 animals. Roller animals expressing GFP were segregated to create the transgenic lines UN1727 (transcriptional reporter) and UN1742 (translational reporter) respectively.

### Creation of spermatheca bag specific RNAi strain

Extra-chromosomal arrays expressing full length *rde-1* under the spermathecal bag specific promoter *fkh-6p* and *fln-1p::GFP*, a spermathecal marker, were UV integrated into the WM27 *rde-1(ne219)* genome. Animals were irradiated with 100 nm UV light for 100 sec and then left to recover for several days. Animals that successfully integrated *fkh-6p::rde-1*; *fln-1p::GFP* were back crossed 3 times with WM27 *rde-1(ne219)*, generating strain UN1524.

### RNA interference

The RNAi protocol was performed essentially as described in Timmons et al (1998). HT115(DE3) bacteria (RNAi bacteria) transformed with a dsRNA construct of interest was grown overnight in Luria Broth (LB) supplemented with 40 μg/ml ampicillin and seeded (150 μl) on NGM plates supplemented with 25 μg/ml carbenicillin and disopropylthio-β-galactoside (IPTG). Seeded plates were left for 24-72 hours at room temperature (RT) to induce dsRNA expression. Empty pPD129.36 vector (“Control RNAi”) was used as a negative control in all RNAi experiments.

Embryos from gravid adults were collected using an alkaline hypochlorite solution as described by Hope (1999) and washed three times in M9 buffer (22 mM KH_2_PO_4_, 42 mM NaHPO_4_, 86 mM NaCl, and 1 mM MgSO_4_) (‘egg prep’). Clean embryos were transferred to supplemented NGM plates seeded with HT115(DE3) bacteria expressing dsRNA of interest and left to incubate at 23°C for 50-56 hours depending on the experiment. In experiments where larvae, rather than embryos, were transferred to RNAi plates (*kin-1(RNAi)* and *kin-2(RNAi))*, adults were ‘egg prepped’ into Control RNAi and left to incubate at 23°C for 32-34 hours, after which they were moved to bacterial lawns expressing the dsRNA of interest and returned to 23°C, and imaged 50-56 after the egg prep.

### Population assay

Embryos collected via an ‘egg prep’ as previously described were plated on supplemented NGM seeded with RNAi bacterial clones of interest. Plates were incubated at 23°C for 54-56 hours or until animals reached adulthood. Upon adulthood nematodes were killed in a drop of 0.08 M sodium azide (NaAz) and mounted on 2% agarose pads to be visualized using a 60x oil-immersion objective with a Nikon Eclipse 80i epifluorescence microscope equipped with a Spot RT3 CCD camera (Diagnostic instruments; Sterling Heights, MI, USA). Animals were scored for the presence or absence of an embryo in the spermatheca as well as entry defects such as gonad arms containing endomitotically duplicating oocytes (Emo) [52]. A Fisher exact *t*-test (two dimensional *x*^*2*^ analysis) using GraphPad Prism statistical software was used to compare the percent of occupied spermathecae between control RNAi and all other RNAi treatments.

### Wide-field Fluorescence Microscopy

#### Acquisition

All Differential Interference Contrast (DIC) and fluorescent images were taken using a 60x oil-immersion objective with Nikon Eclipse fluorescent microscope equipped with a Spot RT3 CCD camera (Diagnostic instruments; Sterling Heights, MI, USA) or Spot RT39M5 with a 0.55x adapter unless otherwise stated. Fluorescence excitation was provided by a Nikon Intensilight C-HGFI 130W mercury lamp and shuttered with a SmartShutter (Sutter Instruments, Novato CA, USA). For acquisition of time-lapse images young adult animals were immobilized with 0.05 micron polystene polybeads diluted 1:2 in water (Polysciences Inc., Warrington, PA, USA) and mounted on slides with 5% agarose pads. Time lapse GCaMP imaging was captured at 1 frame per second, with an exposure time of 75 ms and a gain of 8 for movies obtained with the Spot RT3CCD camera, and with an exposure time of 20 ms and a gain of 8 for movies obtained with the Spot RT39M5. Time-lapse images were only taken of the first 3 ovulations, with preference for the 1^st^ ovulation. The same microscopy image capture parameters were maintained for all imaging.

### Image Processing

All time-lapse GCaMP3 images were acquired as 1600×1200 pixels for the Spot RT3 OCCD camera or 2448×2048 for the RT39M5 camera and saved as 8-bit tagged image and file format (TIFF) files. All image processing was done using a macro on Image J [71]. All time-lapse images were oriented with the sp-ut valve on the right of the frame and registered to minimize any body movement of the paralyzed animal. An 800×400 region of interest for the Spot RT3 OCCD and 942×471 for the RT39M5 camera encompassing the entire spermatheca was utilized to measure the GCaMP3 signal. The average pixel intensity of each frame was calculated using a custom ImageJ macro [5]. Ca^2+^ pixel intensity (F) was normalized to the average pixel intensity of the first 30 frames prior to the start of ovulation (F_0_) and plotted against time. Data analysis and graphing were performed using GraphPad Prism. Kymograms were generated using an ImageJ macro that calculated the average pixel intensity of each column of a frame and condensing it down to one line per frame of the time-lapse image. Every frame of the time-lapse image was stacked on top of each other so to visualize Ca^2+^ dynamics of representative ovulations in both space and time.

### Time Point Metrics

During time-lapse image processing four timepoints are recorded for each ovulation. These include (1) start of oocyte entry: the time when the spermatheca is beginning to be pulled over the incoming oocyte, (2) distal spermathecal closure: the time when the distal spermatheca closes over the oocyte and the oocyte is completely in the spermatheca, (3) sp-ut valve opening: the time when sp-ut valve starts to open and spermathecal exit begins (4) sp-ut valve closure: the time when the sp-ut valve completely closes and the embryo is fully in the uterus. These times were used to calculate both the total transit time, dwell time and exit time of each ovulation. The total transit time is defined as the time from the start of oocyte entry to sp-ut valve closing. Dwell time is defined as the amount of time between the distal neck closing to the sp-ut valve opening. Dwell time captures how long the embryo is in the spermatheca with both the neck and sp-ut valve closed. Exit time is defined as the amount of time from the sp-ut valve opening until the embryo is completely in the uterus and the sp-ut valve closes again behind it.

### Statistics

Either a Fishers exact *t*-test (two dimensional *x*^*2*^ analysis) or a one-way ANOVA with a multiple comparison Tukey’s test were conducted using GraphPad Prism on total, dwell and exit transit times of ovulations acquired via time-lapse imaging. For population assays, statistics were performed using the total number of unoccupied spermathecae compared with the sum all other phenotypes. N is the total number of spermathecae counted. For transit phenotype analysis, statistics were performed using the total number of oocytes that exited the spermatheca successfully compared to the sum of all other phenotypes. Fisher’s exact test were used for both population assays and transit phenotype analysis. Stars designate statistical significance (**** p<0.0001, *** p<0.005, ** p<0.01, * p<0.05).

## Supplemental Figure Legends

**Supplemental Movie 1: Ca**^**2+**^ **oscillations in the spermatheca of a wild type hermaphrodite grown on control RNAi.** A representative wild type ovulation of an animal expressing GCaMP3. Oocyte enters from the right at 30 seconds. There is an initial pulse of Ca^2+^ in the sp-ut valve, followed by Ca^2+^ oscillations that originate from the distal spermatheca on the right, and propagate to the proximal spermatheca on the left until embryo exit.

**Supplemental Movie 2: Ca**^**2+**^ **oscillations in the spermatheca of a wild type hermaphrodite grown on *gsa-1(RNAi).*** A representative *gsa-1(RNAi)* ovulation. Oocyte enters from the right at 30 seconds. The sp-ut valve shows increased Ca^2+^ signal after oocyte entry, and signal remains elevated throughout movie. Embryo is trapped in the spermatheca.

**Supplemental Movie 3: Ca**^**2+**^ **oscillations in the spermatheca of a wild type hermaphrodite grown on *kin-1(RNAi).*** A representative *kin-1(RNAi)* ovulation. Oocyte enters from the right at 30 seconds. The sp-ut valve shows increased Ca^2+^ signal after oocyte entry, and signal remains elevated throughout movie. Additionally, there is increased Ca^2+^ signal only in the proximal spermatheca. Embryo is pushed back into the gonad arm and then re-enters the spermatheca.

**Supplemental Movie 4: Ca**^**2+**^ **oscillations in the spermatheca of a wild type hermaphrodite grown on *kin-2(RNAi).*** A representative *kin-2(RNAi)* ovulation. Oocyte enters from the right at 30 seconds. Pulses of Ca^2+^ propagate from distal to proximal immediately after oocyte entry. The sp-ut valve opens almost immediately. The embryo did not exit during the duration of the movie.

## Acknowledgments

We thank Charlotte Kelley and other members of the Cram and Apfeld laboratories for helpful comments on the manuscript. Some *C. elegans* strains used in this study were provided by the Caenorhabditis Genetics Center, which is funded by the National Center for Research Resources, National Institutes of Health. This work was supported by a grant from the National Institutes of Health National Institute of General Medical Sciences (GM110268) to E.J.C.

